# Life cycle of arabidopsis in the international space station - Growth direction of the inflorescence stems in the presence of light under microgravity

**DOI:** 10.1101/2020.09.30.320051

**Authors:** Umi Yashiro, Ichirou Karahara, Sachiko Yano, Daisuke Tamaoki, Fumiaki Tanigaki, Toru Shimazu, Daisuke Masuda, Haruo Kasahara, Seiichiro Kamisaka

## Abstract

In the “Space Seed” experiment performed in Kibo module of the International Space Station, growth direction of the inflorescence stem of arabidopsis was examined under space 1 *G*, μ*G*, and ground 1 *G* conditions in the presence of light. The stems grew almost upright (vertical to the surface of seedbed) under ground 1 *G*. Although the stems were primarily upright both under space 1 *G* and μ*G*, they tilted slightly. The tilting of the stems under space 1 *G* was indicated to be due to tilting of the artificial gravitational acceleration vectors produced on the centrifuge. The tilting of the stems under μ*G* was suggested to be due to the pressure of directional airflow produced by ventilation.

## 1. INTRODUCTION

Among various environmental factors, gravity and light play a crucial role in regulating growth direction of plant shoots. In the absence of gravity, light has been considered to be able to mimic the effect of gravity on growth direction of plant shoots. It has been shown that plant shoots have grown toward light source under microgravity (μ*G*) conditions [1] [2]. However, the growth direction of the stem has not yet examined precisely under μ*G* in the light. It is important to understand how form of plants is regulated in the absence of gravity for designing efficient plant cultivation systems in space including light conditions. In the “Space Seed” experiment [3], a centrifuge has been installed to produce artificial gravity, which enables to investigate effects of gravity acceleration by comparing precisely between μ*G* and 1 *G* on orbit. In the present study, we attempted to determine the precise direction of shoot growth under μ*G* conditions in the light.

## 2. MATERIALS AND METHODS

The “Space Seed” experiment was performed on board KIBO module from 2009 to 2010. Seeds of *Arabidopsis thaliana* (L.) Heynh. ecotype Columbia-0 were sterilized and glued to surface of a rockwool slab, which was placed in a plant growth chamber made of transparent polycarbonate (H x W x D = 60 x 60 x 50 mm, outer size). The chamber was installed in a plant growth apparatus called Plant Experiment Unit (PEU) [3]. An LED plate was placed on the chamber for continuous illumination (red and blue radiation in a ratio of 3:1) at the intensity of 110 μmol m^-2^s^-1^ at the surface of stainless plate placed on the surface of seedbed. A CCD camera was used to obtain images of the inside of the chamber. PEUs were installed in an incubator called the Cell Biology Experiment Facility (CBEF) [3] onboard the Kibo module. The CBEF was equipped with a centrifuge generating artificial gravity to perform 1-*G* control experiment as well as microgravity experiment on board. A ground control experiment was simultaneously performed in Toyama, Japan. Digital photographs were taken and downlinked on a daily basis to monitor growth of arabidopsis plants. Using digital images, the growth directions of inflorescence stems were assessed by analyzing tilt angles between each inflorescence stem and the vertical line, which was perpendicular to the surface of the stainless plate. The growth direction was determined before each inflorescence stem reached ceiling of a growth chamber. Positive and negative angles represent clockwise and counterclockwise inclination of the stems from the vertical line, respectively.

## 3. RESULTS AND DISCUSSION

Judging from the downlinked images, inflorescence stems grew primarily upward under μ*G* in space. This indicates that phototropism replaces gravitropism in the absence of gravity. When the growth directions of inflorescence stems were carefully observed, however, they appeared to be tilted slightly in the clockwise direction both under space 1 *G* and μ*G* when compared to Toyama (ground) 1 *G*. Precise tilting angles of inflorescence stems were examined (Fig. 1). The tilt angle of the stem was - 0.1 ± 0.4 degree (mean ± SE, n=45) under Toyama 1 *G*, while that was - 3.9 ± 0.8 degree (n=35) under space 1 *G*. Interestingly, negative angle of - 3.0 ± 1.0 degree (n=38) was observed also under μ*G*. When the data were pooled, a significant difference was found in the angles between Toyama 1 *G* and space 1 *G* (Kruskal-Wallis test, *P* <0.001).

**Fig. 1.**
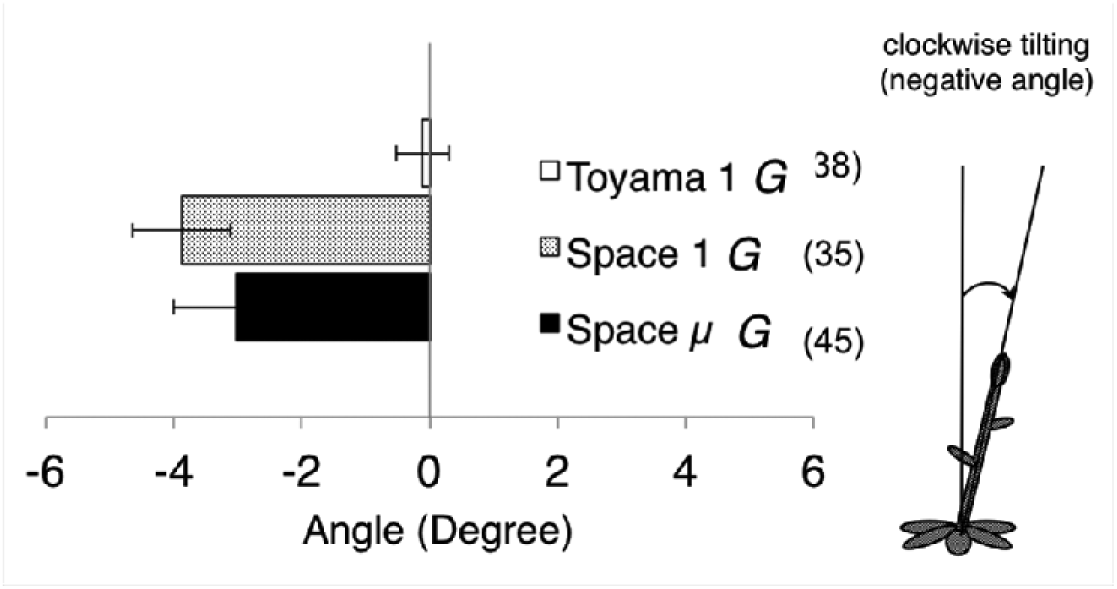
Effects of microgravity on the mean tilt angle of the inflorescence stem of arabidopsis in the light. Values are mean ± SE. Numbers of samples (n) are shown in parenthesis. Negative angles represent clockwise tilting of the stems from the vertical line. Values for mean, SE, and n were calculated across all plants

The area of seedbed was divided into three subareas (left, center, right) and distribution of tilt angles of inflorescence stems was examined. The stems tilted more in the clockwise direction in the left subarea compared to the other subareas both under space 1 *G* and μ*G* (Fig. 2). The overall distribution of space 1 *G* and μ*G* appeared broader than Toyama 1 *G*. Actually, significant differences were found in the variance between Toyama 1 *G* and space 1 *G* or μ*G* (*F*-test, two-tailed, *P* <0.01) when the data were pooled.

**Fig. 2.**
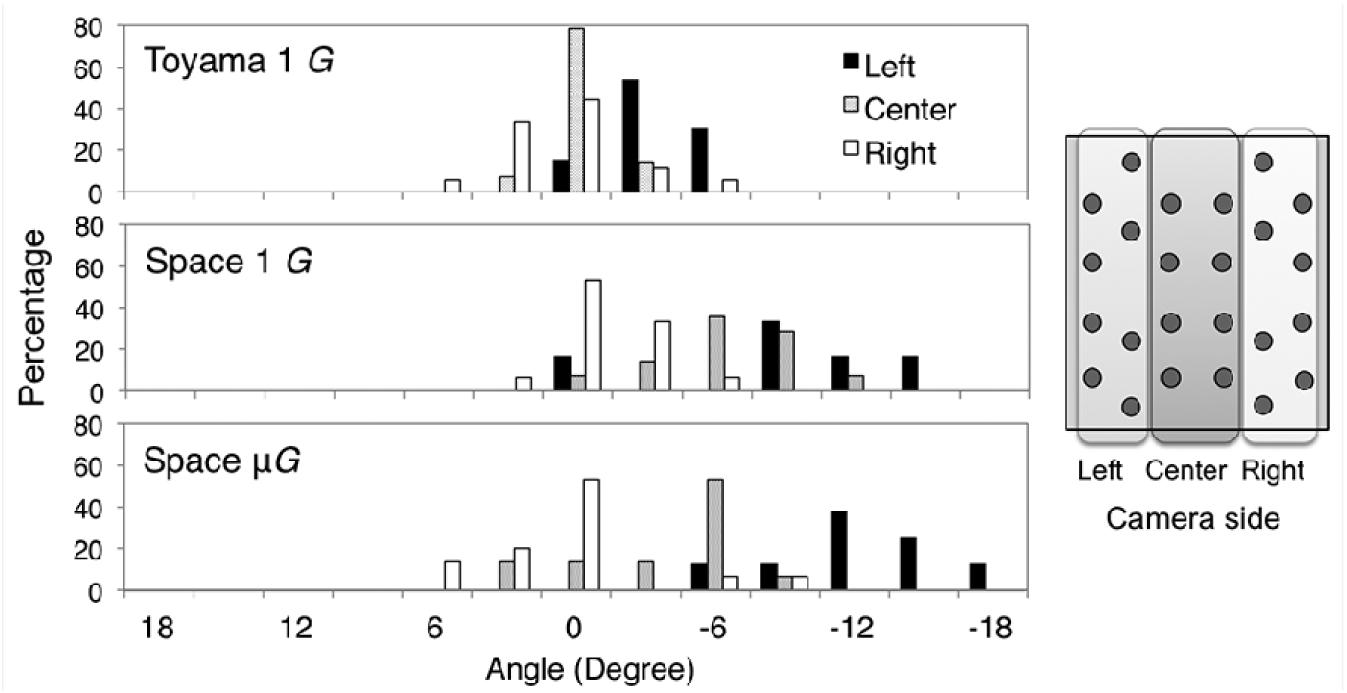
Effects of microgravity on the distribution of tilt angle of inflorescence stems of arabidopsis. Surface of a seedbed was divided into three subareas (left, center, right). The schematic illustration shows positions of each plant and the subareas viewed from the top of a chamber

Both under space 1 *G* and μ*G*, the tilt angle of the stem became smaller (more clockwise) when the position was closer to the left side. As a matter of fact, the direction and the magnitude of gravitational acceleration vector was different at every location in a chamber because the artificial gravitational acceleration was produced using a centrifuge (Fig. 3). Tilt angle of the vector from the vertical line became larger when the position was closer to the left side. Therefore, it is possible that the tilting of the stem observed in space 1 *G* was due to the tilting of the artificial gravitational acceleration vectors produced in PEUs on the centrifuge.

**Fig. 3.**
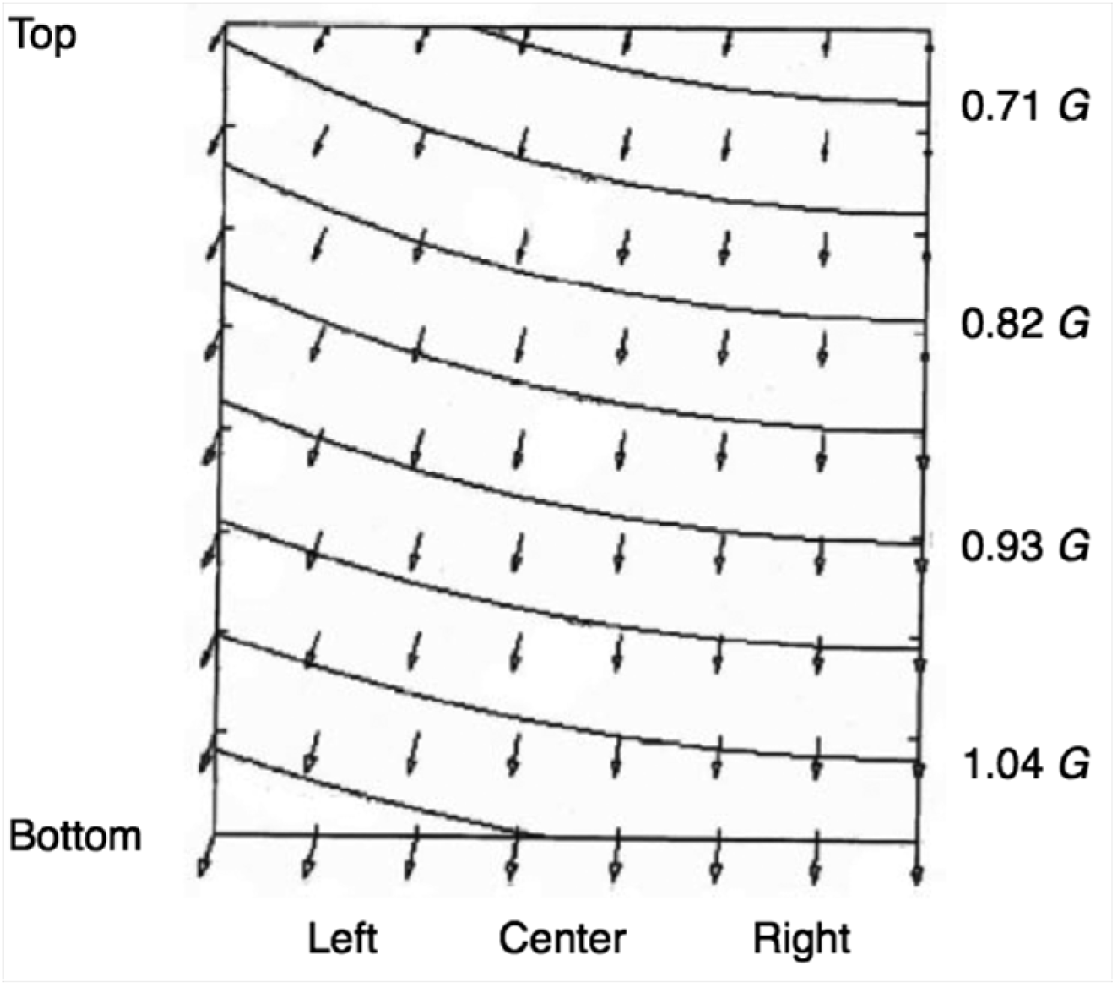
A schematic diagram showing the direction and the magnitude of gravity acceleration vectors (arrowheads) in a chamber of space 1 G viewed from the camera side of a chamber

On the other hand, negative tilt angle of the stem was also observed under μ*G*. This cannot be explained by the artificial gravitational acceleration. One of possible explanations for this negative tilting angle may be the effect of the pressure of directional air flow caused by ventilation. Air inlets and outlets for ventilation were located along the upper part of the left sidewall and the lower part of the right side wall of a chamber, respectively. Therefore, air flowed from the upper left side to the lower right side. This directional flow of air occurred under each gravity condition. Slight clockwise tilting of the stem was observed in the left subarea even under Toyama 1 *G*, which is possibly due to this directional airflow. Therefore, the direction of the growth (clockwise tilting) is possibly caused by the pressure of this directional airflow also under space 1 *G*.

The fact that the overall tilt angle of the stem was close to zero degree under Toyama 1 *G* indicates that gravitropism masks the effect of directional airflow in the present experimental condition. The tilting of the stem under space 1 *G* was indicated to be primarily due to the tilting of the artificial gravitational acceleration vectors. Under μ*G*, the direction of the stem growth should primarily be determined by light. However, the pressure of directional airflow produced by ventilation is suggested to affect substantially the growth direction of the stem.

## 4. ACKNOWLEDGEMENTS

The present work was supported by JSPS KAKENHI Grant (Nos. 21570064 and 24620003) to I. K..

